# MoMo: Discovery of post-translational modification motifs

**DOI:** 10.1101/153882

**Authors:** Alice Cheng, Charles E. Grant, Timothy L. Bailey, William Stafford Noble

## Abstract

**Motivation:** Post-translational modifications (PTMs) of proteins are associated with many significant biological functions and can be identified in high throughput using tandem mass spectrometry. Many PTMs are associated with short sequence patterns called “motifs” that help localize the modifying enzyme. Accordingly, many algorithms have been designed to identify these motifs from mass spectrometry data.

**Results:** MoMo is a software tool for identifying motifs among sets of PTMs. The program re-implements two previously described algorithms, Motif-X and MoDL, packaging them in a web-accessible user interface. In addition to reading sequence files in FASTA format, MoMo is capable of directly parsing output files produced by commonly used mass spectrometry search engines. The resulting motifs are presented to the user in an HTML summary with motif logos and linked text files in MEME motif format.

**Availability:** Source code and web server available at http://meme-suite.org

**Supplementary information:** Supplementary figures are available at *Bioinformatics* online.

## 1 Introduction

Protein post-translational modifications (PTMs) control or otherwise participate in a wide range of biological activities, including critical regulatory functions. Dysregulation of some types of modifications, such as phosphorylation, has been implicated in a variety of diseases. The best characterized modifications include methylation, acetylation, phosphorylation, and sumoylation, but hundreds of less common types of modifications have been identified and catalogued in various databases. Recently, high-throughput mass spectrometry has greatly increased our ability to identify PTM sites.

In parallel with these experimental efforts is an ongoing stream of research devoted to characterizing PTM sites *in silico*. The PTM motif discovery problem differs from traditional sequence motif discovery in four important ways: (1) the candidate motif sites can be aligned relative to the (known) PTM, (2) the number of identified sites can be quite large, (3) the motifs tend to have low information content, and (4) the set of identified sites is usually contaminated with false positive identifications. Note that, like traditional motif discovery, the data set may contain a mixture of motifs, each corresponding to different modifications. The latter is particularly relevant to phosphorylation, where different types of kinases phosphorylate variant sequence motifs.

We have developed a software tool, MoMo (for “modification motifs”), to allow MEME Suite users to identify protein PTM motifs. MoMo re-implements two existing modification motif discovery methods. Motif-X (Schwartz and Gygi, 2005) was one of the first such algorithms and is by far the most widely used method for PTM motif analysis: according to Google Scholar, the Motif-X publication has 602 citations, whereas the publications associated with the other six algorithms in Table 1 have a total count of 125 citations. The second algorithm in MoMo is the “Motif Description Length” (MoDL) algorithm (Ritz *et al.*, 2009). Although MoDL is not yet as widely used as Motif-X, we chose to reimplement it because MoDL employs a much more rigorous approach to the PTM motif discovery problem, formulating a clear objective and proposing a greedy heuristic to optimize that objective. Also, these two algorithms are complementary, in the sense that they employ different underlying motif representations: internally, Motif-X employs a “consensus plus mismatch” model in which each position in the motif is represented as a single amino acid or a wildcard character; MoDL, in contrast, uses a regular expression representation, wherein multiple different amino acids are allowed to occur at any given position. In practice, in both algorithms, sequences that match the underlying motif model are summarized in a position-specific scoring matrix that is provided to the user.

**Table 1.**
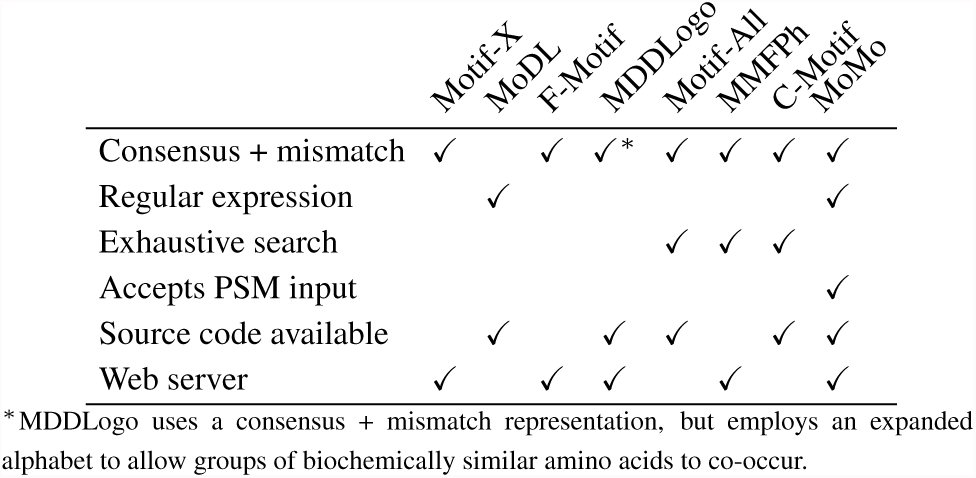
Comparison of PTM motif discovery tools.

We also produced a web portal for MoMo, integrated into the existing MEME Suite portal. This web page provides various user-level parameters for controlling the type of motifs found and provides an option for selecting which algorithm to use. The server provides an interactive JavaScript interface, allowing users to upload large PTM sequence sets for analysis. Importantly, to enhance usability, the portal works directly with database search results files produced by popular mass spectrometry search engines, including Comet, MS-GF+, Tide, etc. (reviewed by (Nesvizhskii, 2010)) Results are provided in a user-friendly HTML format, similar to other MEME Suite outputs. By providing source code and a web server, and supporting two complementary algorithms, MoMo provides broader functionality than existing tools (Table 1).

## 2 Implementation

MoMo is implemented in C and is available as a command line tool and through the web interface.

MoMo takes as input either a tab-delimited peptide-spectrum match (PSM) file or a set of fixed-width sequences centered around the site of modification. The latter can be in either prealigned format, where each sequence is separated by a newline, or FASTA format. After uploading the modification dataset, the user is given the option of specifying the protein database that was used in the search. The user can provide their own protein database (in FASTA format), select a database from a drop-down menu, or choose not to use any protein database at all. A protein database is required when searching for motifs using the Motif-X and MoDL algorithms but is not required when creating simple motifs based on the frequency of each residue surrounding the modification. When given both a tab-delimited PSM file and a protein database, the software converts the PSM file into a set of fixed-width sequences by searching the protein database for an occurence of the peptide and then extracting the flanking residues. A set of background sequences is constructed by finding, for each modified residue in the foreground set, an occurrence of that residue in the protein database. Finally, the user has the option of identifying motifs separately for each unique modification mass, or for each unique residue+mass combination.

MoMo produces as output an HTML report, with each motif in Portable Network Graphics (.png) format and including links to the motifs in the MEME plain text motif format. The MoMo web server allows the user to upload one or more PSM files and searches them jointly. Search results are stored online, and the user is notified of their availability via email.

## 3 Results

To validate MoMo, we used a phosphorylation dataset consisting of 305 PTM sequences from the Phospho.ELM database (Dinkel *et al.*, 2010). We first verified that MoMo produces the same motifs as the original implementations of Motif-X and MoDL (Supplementary Note 1). Next, we used the known kinase family assignments from Phospho.ELM to evaluate the quality of the motifs discovered by various algorithms, computing the adjusted Rand index between the true clusters and the clusters induced by the discovered motifs (details in Supplementary Note 2). The results (Figure 1) show that none of the methods succeeds in achieving a clustering that is very similar to the best clustering possible using position-specific scoring matrices (marked “Ideal” in the figure). MDDLogo performs well but can only find two clusters. MoDL performs well but can only find three clusters, and Motif-X starts to perform well only when the number of discovered clusters is larger than the number of true clusters.

**Fig. 1.**
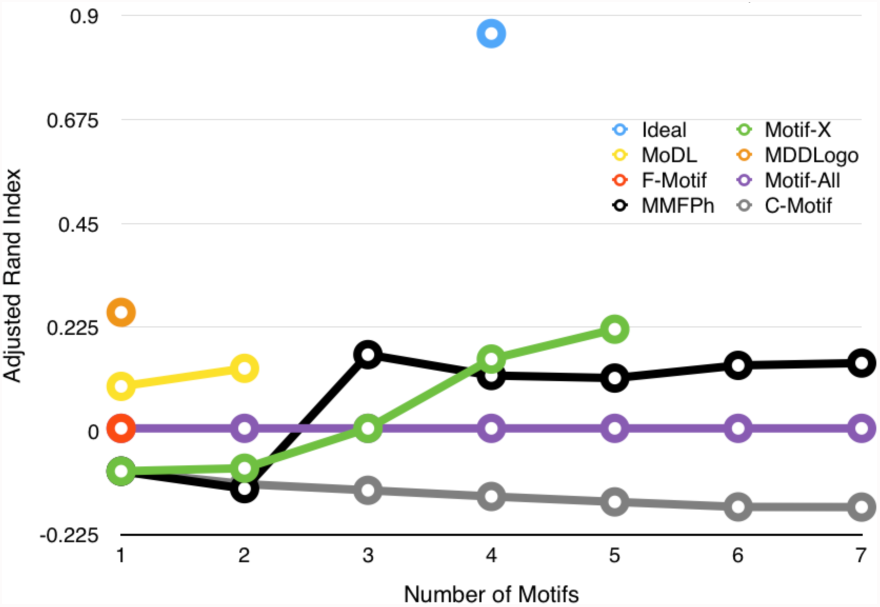
Comparison of PTM motif discovery algorithms relative to a Phospho.ELM benchmark. The figure plots, for each algorithm, the adjusted Rand index between the clustering induced by the discovered motifs relative to the “true” clustering provided by Phosho.ELM. The “Ideal” clustering is constructed by scanning with motifs built from the ground truth clusters. The Rand index has a maximum value of 1 and an expected value of 0 for a random clustering.

Of course, this small benchmark experiment is not conclusive. Our goal is primarily to demonstrate the validity of our implementation, and to suggest that further work is still needed to compare and contrast various PTM motif discovery approaches on realistic benchmark data sets. We hope that MoMo will be a useful tool, both in the context of such evaluations and in the analysis of new PTM data sets.

